# Multicenter comparison of the Cobas 6800 system with the RealStar RT-PCR kit for the detection of SARS-CoV-2

**DOI:** 10.1101/2020.06.29.179184

**Authors:** Marc Wirden, Linda Feghoul, Mélanie Bertine, Marie-Laure Nere, Quentin Le Hingrat, Basma Abdi, David Boutolleau, Valentine Marie Ferre, Aude Jary, Constance Delaugerre, Anne-Genevieve Marcelin, Diane Descamps, Jérôme Legoff, Benoit Visseaux, Marie-Laure Chaix

**Affiliations:** Sorbonne Université, INSERM, Institut Pierre Louis d’Epidémiologie et de Santé Publique IPLESP, AP-HP, Hôpital Pitié-Salpêtrière, Laboratoire de virologie, Paris, France; Université de Paris, Département des Agents Infectieux, Service de Virologie, Hôpital Saint-Louis, Paris, France; INSERM UMR 976, Université de Paris, Paris, France; Université de Paris, Assistance Publique – Hôpitaux de Paris, Service de virologie, Hôpital Bichat, Paris, France; UMR 1137-IAME, DeSCID: Decision SCiences in Infectious Diseases control and care, INSERM, Université de Paris, Paris, France; INSERM UMR 944, Université de Paris, Paris, France

**Keywords:** COVID-19, SARS-CoV-2, RT-PCR

## Abstract

**Background:** RT-PCR testing is crucial in the diagnostic of SARS-CoV-2 infection. The use of reliable and comparable PCR assays is a cornerstone to allow use of different PCR assays depending on the local equipment. In this work, we provide a comparison of the Cobas® (Roche) and the RealStar® assay (Altona).

**Methods:** Assessment of the two assays was performed prospectively in three reference Parisians hospitals, using 170 clinical samples. They were tested with the Cobas® assay, selected to obtain a distribution of cycle threshold (Ct) as large as possible, and tested with the RealStar assay with three largely available extraction platforms: QIAsymphony (Qiagen), MagNAPure (Roche) and NucliSENS-easyMag (BioMérieux).

**Results:** Overall, the agreement (positive for at least one gene) was 76%. This rate differed considerably depending on the Cobas Ct values for gene E: below 35 (n = 91), the concordance was 99%. Regarding the positive Ct values, linear regression analysis showed a determination correlation (R^2^) of 0.88 and the Deming regression line revealed a strong correlation with a slope of 1.023 and an intercept of -3.9. Bland-Altman analysis showed that the mean difference (Cobas® minus RealStar®) was + 3.3 Ct, with a SD of + 2.3 Ct.

**Conclusions:** In this comparison, both RealStar® and Cobas® assays provided comparable qualitative results and a high correlation when both tests were positive. Discrepancies exist after 35 Ct and varied depending on the extraction system used for the RealStar® assay, probably due to a low viral load close to the detection limit of both assays.

## INTRODUCTION

The SARS-CoV-2 is the new coronavirus, a member of the subgenus Sarbecovirus (beta-Coronavirus lineage B), responsible for the ongoing pandemic of infectious respiratory disease called COVID-19 (1, 2). This epidemic, declared a Public Health Emergency of International Concern on 30 January 2020 by the World Health Organization (WHO), has spread rapidly around the world and has caused many infections and deaths worldwide (3). To control the outbreaks, many countries have implemented confinement instructions and have closed places of groupings with heavy economic consequences. As recommended by WHO, diagnostic tests by reverse transcription-polymerase chain reaction (RT-PCR) via respiratory samples, should be performed widely to detect infected patients, to follow the pandemic evolution and to help stopping the spread of the clusters (4). RT-PCR testing is also a cornerstone to allow a gradual deconfinement in good sanitary conditions and early detect any viral resurgence. To meet the high demand for these tests and to face the supply difficulties worldwide, the laboratories had to adapt using the different systems available depending on the PCR and nucleic acid extraction equipment already present in their establishment (5–10). Manufacturer-independent evaluation data are still scarce. These tests can be a single-use cartridge, reagent kits for batch testing used with different instruments for the extraction and amplification stages, or fully automated molecular testing platforms. These are real-time RT-PCR which target two or three different regions of the SARS-CoV-2 genome and provide a cycle threshold (Ct) value inversely proportional to the amount of virus. Pre-analytical processing of respiratory samples can be also used to neutralize the virus before testing, and the sample input volume used varies depending on the test performed. All these differences in the pre-analytical and analytical process can have an impact on the sensitivity of the test and the concordance of their results has to be evaluated.

In this study, we compared two different widely used tests in three major Parisian university hospital laboratories. These are the RealStar® SARS-CoV-2 RT-PCR Kit 1.0 (Altona diagnostics, France) which can be associated to different extraction and amplification devices, and the Cobas® SARS-CoV-2 kit used on the Cobas® 6800 system (Cobas 6800; Roche Diagnostics, Mannheim, Germany).

## METHODS

### Samples

In April 2020, 140 patients were included in this prospective study performed in 3 virological laboratories located in Paris (Saint Louis hospital (n=45), Bichat hospital (n=49) and La Pitié-Salpêtrière hospital (n=46)). Then, each laboratory selected 45 to 49 samples firstly detected using the Cobas 6800 with a stratification according to the Ct of the E gene Cobas results, allowing to cover the whole linear range of the assays. Thus, three categories were retained: Ct < 25, Ct between 25 and 34 and with a Ct ≥ 35. Rapidly, in the same day or within 48 hours, the leftover samples stored at +4°C were tested with the RealStar assay. Thirty nasopharyngeal swab samples collected in 2019 (in the pre-epidemic Covid 19 period) were also tested with both techniques (10 in each laboratory).

### Cobas 6800 SARS-CoV-2 test

The Cobas^®^ SARS-CoV-2 test is a single-well dual target assay, which targets the non-structural ORF1a/b region specific of SARS-CoV-2 and the structural protein envelope E gene for pan-sarbecovirus detection. The test used RNA internal control for sample extraction and PCR amplification process control. To take into account the available sample volume and the security conditions required for this virus before loading on the Cobas 6800 system, the pre-analytical protocol has been adapted as recommended by the manufacturer as follows: 400 μl of each sample were transferred at room temperature into barcoded secondary tubes containing 400 μl of Cobas lysis buffer for the SARS-CoV-2 neutralization. Then, the tube was loaded on the Cobas 6800 where 400 μl from those 800 μl were used for RNA extraction, and eluted in 50 μl of which 27 µl were used for the RT-PCR. The test was performed in batches, including one negative and positive control each. According to the manufacturer’s instructions, a tested sample was considered SARS-CoV-2 positive if Cobas 6800 showed positive results either for both ORF1a/b and E genes or for the ORF1a/b gene only. In the case of single E gene positivity, the result should be reported as SARS-CoV-2 presumptive positive and repeated, but were considered as positive for this study.

### RealStar SARS-CoV-2 RT-PCR

The RealStar® SARS-CoV-2 RT-PCR Kit 1.0 assay targets the E gene specific for sarbecoviruses, and the S gene specific for SARS-CoV-2. It includes a heterologous amplification system (Internal Control) to identify possible RT-PCR inhibition and to confirm the integrity of the kit reagents. This kit contains only reagents for the SARS-CoV-2 real-time RT-PCR step, extraction and amplification can be performed with various equipment listed in the kit insert. In this study, RNA extraction was performed with MagNA Pure LC 2.0 System (Roche) (Bichat hospital), QIAsymphony (Qiagen) (Saint Louis hospital) and NucliSENS® EasyMag® (bioMérieux) (Saint Louis hospital and Pitié Salpêtrière hospital) according to manufacturer’s protocol. In each cases, 200 µl of nasopharyngal samples were diluted in 2 ml of lysis buffer and eluted in 50 µl. Ten µl of extracted RNA was used to perform the real-time RT-PCR with the LightCycler^®^ 480 (Roche) in Pitié Salpêtrière hospital or the ABI Prism^®^ 7500 SDS (Applied Biosystems) in the two other laboratories. All these instruments are listed into the RealStar assay instructions. The sample was considered as positive if at least one of both targets was detected.

### Statistical analysis

Statistical analyses were performed on GraphPad Prism version 6.0. The negative results obtained with the RealStar test were excluded from the analyses. All tests were two-sided, with p values of <0.05 denoting statistical significance. The Ct values obtained with both assays were compared in Wilcoxon tests, and we presented correlation curves with the coefficient of determination, R^2^. Bland-Altman analysis was used to represent the degree of agreement between the Cobas 6800 System and the RealStar^®^ SARS-CoV-2 RT-PCR based on the mean difference and standard deviation (SD) of the positive results. The comparison between the EasyMag and QIAsymphony extraction was done with a paired-samples Student test.

## RESULTS

### Comparison of the Cobas® 6800 System with the RealStar® kit

A total of 170 patient samples were included in this study: 30 collected in 2019, before the French epidemic period, and 140 with a positive detection for SARS-CoV-2 with the Cobas 6800. All the 30 samples collected in 2019 before the epidemic period were negative with both Cobas 6800 and RealStar assays. The qualitative results of the 140 selected samples are summarized in Table 1. Overall, the agreement (positive with the two tests regardless of the gene detected) was 76%. Of note, 3 samples positive in gene E with the COBAS 6800 were negative in gene E but positive in gene S with the RealStar assay. However, this rate differed considerably depending on the Cobas 6800 E Ct: below 35 (n = 91), the concordance was 99%. Only one sample with a Cobas 6800 E Ct at 34.3 was negative in RealStar assay with an EasyMag extraction. For samples with a Cobas 6800 E Ct ≥ 35 (n = 49) only 14/49 was positive in both techniques with a concordance of 29%.

**Table 1:** Agreement between the Cobas 6800 SARS Cov-2 and the RealStar^®^ SARS Cov-2 results according to each gene. E: envelope, S: spike, ORF: open reading frame

|  | gene | Cobas 6800 |  | total |
| --- | --- | --- | --- | --- |
|  |  | E+ ORF-1+ | E+ ORF-1- |  |
| **RealStar<sup>®</sup>** | E+ S+ | 95 (67.9%) | 5 (3.6%) | 100 (71.4%) |
|  | E+ S- | 2 (1.4%) | 2 (1.4%) | 4 (2.9%) |
|  | E- S+ | 2 (1.4%) | 1 (1.4%) | 3 (2.1%) |
|  | E- S- | 16 (11.4%) | 17 (12.1%) | 33 (23.6%) |
| **total** |  | 114 (81.4%) | 25 (18.5%) | 140 (100%) |

For the gene E Ct < 35 obtained with Cobas® 6800, the median of the value obtained with RealStar® assay was 23.5, 23.4 and 18.6 with EasyMag, QIAsymphony and MagNA Pure, respectively. Moreover, for samples with a Ct ≥ 35 with Cobas 6800 (n=49), the detection rate observed with the RealStar assay differed depending on the extraction system, 1/13 with EasyMag, 1/15 with the QIAsymphony and 13/21 with MagNA Pure.

Regarding the positive Ct values of gene E (n=104), linear regression analysis revealed a R^2^ of 0.88 and the Deming regression line revealed a strong correlation with a slope of 1.023 and an intercept of -3.9 (Fig. 1a). The Bland Altman plot showed higher Ct values for the Cobas 6800 with a homogeneous distribution up to Ct 35 with a mean difference (Cobas 6800 minus RealStar) of + 3.3 Ct and a SD of + 2.3 Ct (Fig. 1b).

**Figure 1.**
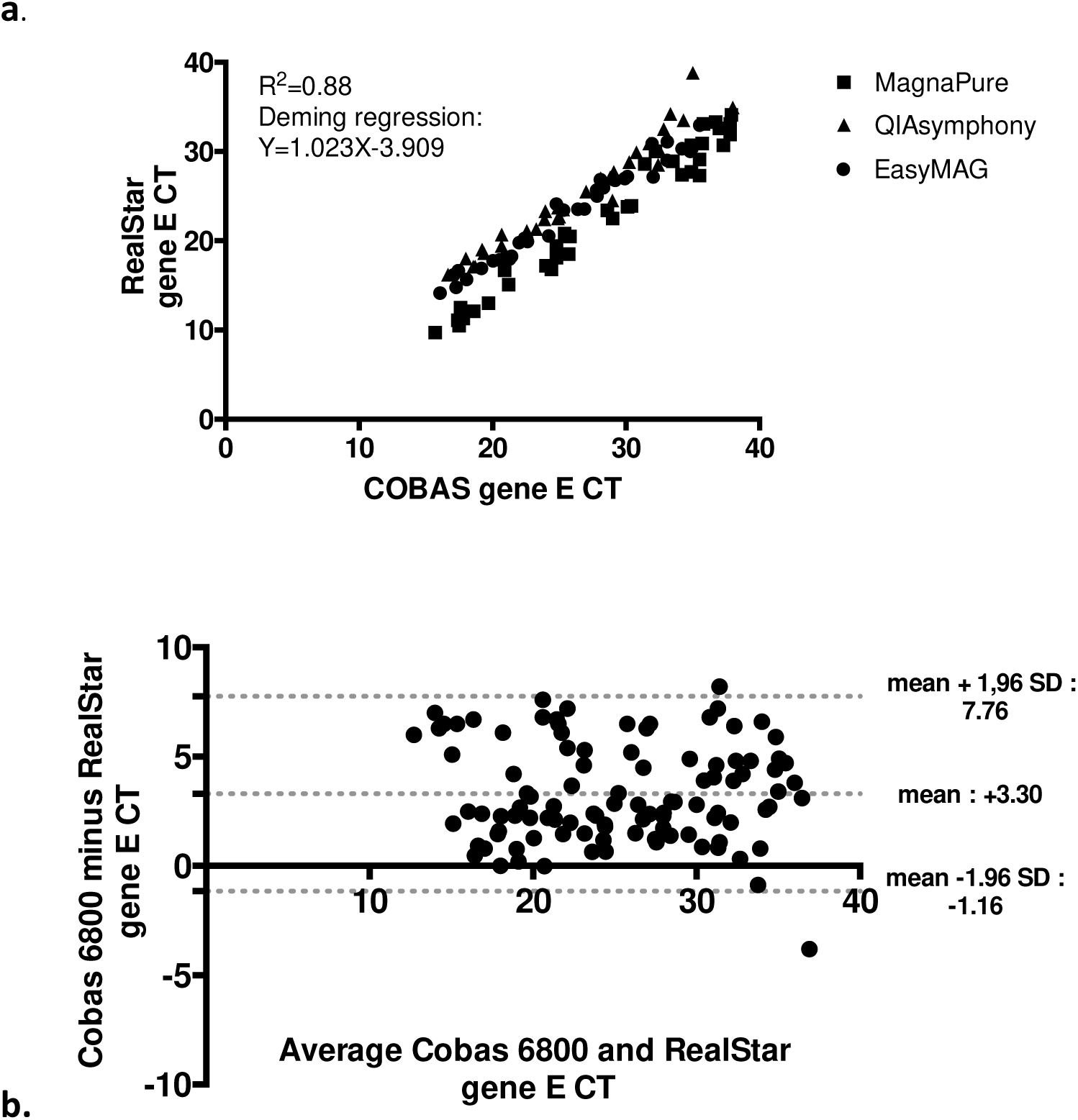
Correlation Curve (a) and Bland-Altman plot (b) for the Ct values of gene E with Cobas 6800 versus RealStar, when both assays are positive (n =104). For the correlation curve the samples extracted by MagnaPure are represented by squares, those with QIAsymphony by triangles and those by EasyMag by dots.

### Comparison of the RealStar results after extraction with EasyMag and QIAsymphony systems

In Saint Louis hospital, 45 samples previously detected with the Cobas 6800 (15 with a Ct < 25, 15 with a Ct between 25 and 34 and 15 with a Ct ≥ 35) were extracted both with the EasyMag kit and the QIAsymphony kit before RealStar testing. All the samples with a Ct < 35 (n=30) were positive regardless of the extraction system. Among the 15 samples with a Cobas 6800 Ct ≥ 35, all were negative after EasyMag extraction while 2 samples were positive after QIAsymphony extraction (Ct: 34.9 and Ct: 38.8). We found a R^2^ of 0.99 and the Deming regression revealed a strong correlation with a slope of 0.99 and an intercept of -0.81 (Fig. 2a). Bland-Altman analysis showed that the mean difference (QIAsymphony minus EasyMag) was + 1.1 Ct, with a SD of - 0.70 Ct (Fig. 2b). Two differences exceeded 5 Ct corresponding to the 2 samples positive using QIAsymphony and negative with EasyMag. Although there was no significant difference in Ct values for the gene E (p=0.21), we have a significant difference in Ct values for the gene S (p<0.0001, mean Ct gene S=1.19, 95% CI: 1.95 to 1.63) in favor of EasyMag.

**Figure 2.**
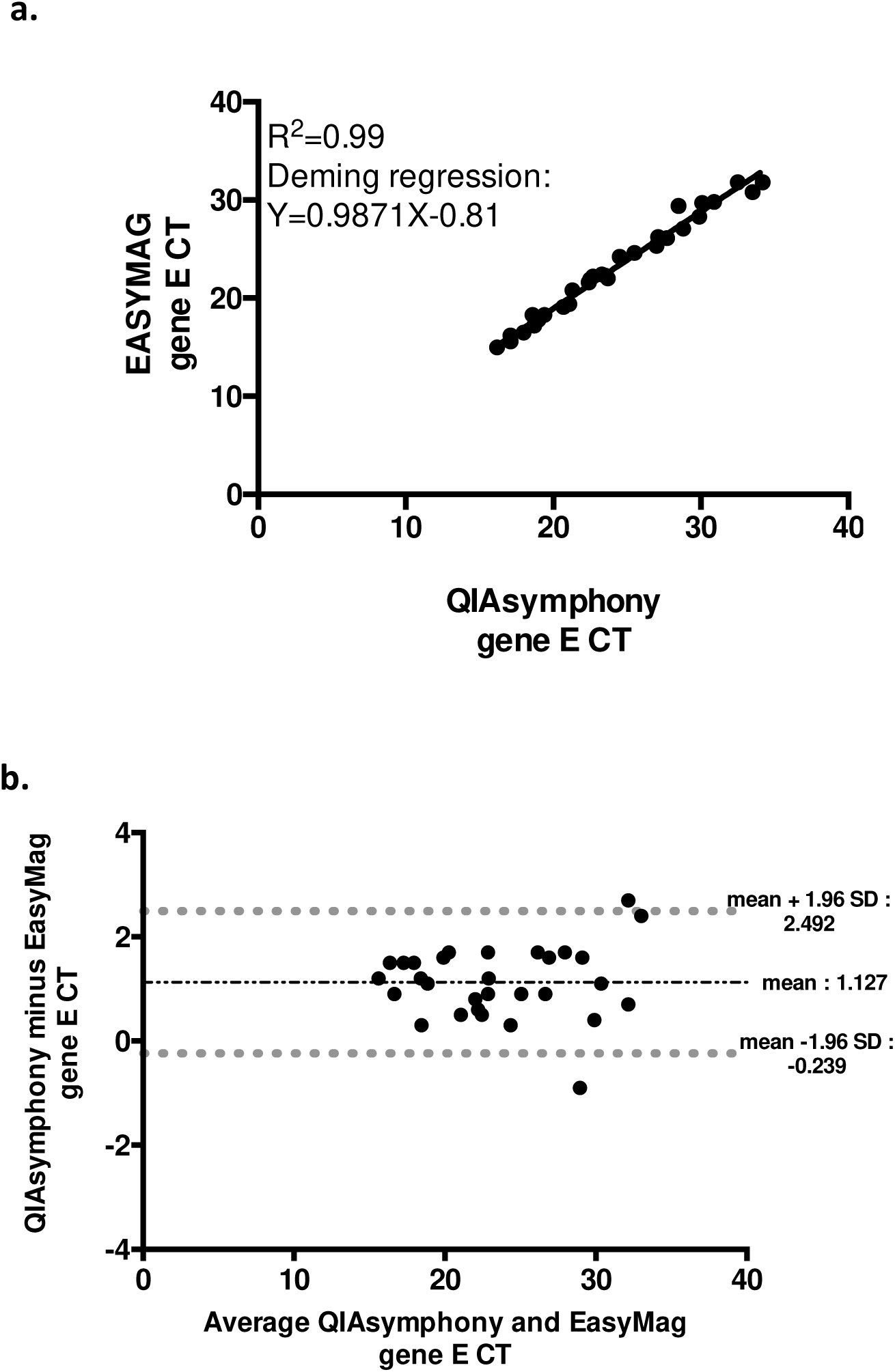
Correlation Curve (a) and Bland Altman plot (b) for the Ct values of gene E RealStar for QIASymphony versus EasyMag extraction, when both assays are positive (n =30).

### Comparison of the detection of gene E and ORF for Cobas 6800

Amplification of gene E (139/140, 99.3%) was more frequently positive compared to amplification of gene ORF (115/140, 82.1%). A R^2^ of 0.84 and the Deming regression line revealed a good correlation with a slope of 1.22 and an intercept of -6.3.

### Comparison of the detection of gene E and S for RealStar® RT-PCR

Amplification of gene E (104/140, 74.3%) and amplification of gene S (115/140, 82.1%) seem to be equivalent. A R^2^ of 0.93 and the Deming regression line revealed a strong correlation with a slope of 1.01 and an intercept of -0.4.

## DISCUSSION

In this study, two very different assays were compared: the RealStar assay used with various extraction equipment, allowing a use in a wide range of PCR laboratories, and the Cobas 6800 kit used with the fully automated Cobas 6800 platform, allowing more intensive workflows. As all other SARS-CoV-2 PCR diagnostic tests, both assays are qualitative but yield a Ct value inversely proportional to the amount of virus. In our work, below a Cobas 6800 Ct value of 35, the qualitative results are highly concordant and the Ct values have a high correlation even though the values of RealStar are lower than those of the Cobas 6800. Above a Cobas 6800 value Ct of 35, the RealStar failed to detect about one third of the SARS-CoV-2 genes while COBAS 6800 detected at least one of both targets. However, this observation is impacted by the extraction method in use, as demonstrated by the slightly lower Ct values and higher positivity rate observed with the MagNA Pure system among samples showing E gene Ct ≥35 with the COBAS 6800. This suggests a better extraction process with the MagNA Pure system. The comparison, from same samples, between EasyMag and QIAsymphony systems showed a slight improvement for the SARS-CoV-2 detection with QIAsymphony. Among samples with Ct values above 35, the E target is mostly the only gene detected with the Cobas 6800 assay. This is in accordance with the Cobas 6800 insert information reporting a higher sensitivity for the E gene detection than for the ORF1/a, and also a drop in the positivity rate above 35 Ct for the E target. This may explain why the RealStar test yielded many negative results in such cases as both tests probably reached their detection limits. This is a limitation of our study as we did not assessed comparatively the limit of detection of the two methods but the reliability of their Ct values among COBAS 6800 positive samples, excluding those that could be negative with COBAS 6800 and positive with RealStar in this range of low viral loads. Our work highlights the impact of the extraction system on the sensitivity of the RealStar assay.

Overall, we demonstrated the good performances and concordance between the two assays, at least for viral loads above the detection limit of both assays. This concordance allows to reliably compare Ct values obtained from both methods. However, the variations observed between the Ct values of the two assays, evaluated here as about 3.5 additional Ct with the Cobas 6800 assay, has to be taken into account for Ct values follow-up done for the most severe patients in case of successive use of the two methods, depending of reagent and analyser availability.

## Acknowledgments

We acknowledge all the laboratory staff of Saint Louis hospital virology department, Bichat hospital virology department and La Pitié-Salpêtrière hospital virology department.

